# Persistence of commensal *Escherichia coli* strains and plasmid transfer in a healthy adult

**DOI:** 10.64898/2026.11.09.765220

**Authors:** Robert A. Moran, Ruth M. Hall

## Abstract

Little is known about the ecological dynamics of *Escherichia coli* strains in the healthy human intestinal tract. Here, the composition of commensal *E. coli* in a healthy adult was tracked over 700 days. 100 *E. coli* isolates recovered directly from each of 21 faecal samples collected at monthly intervals were characterised. In total 15 strains were recovered; five were resistant to one or more antibiotics. Five ‘persistent’ strains were found in multiple samples spanning 1- to 5-month periods. The relative abundance of persistent strains ranged from 1-100% of *E. coli* per sample, but they were dominant (>80%) in 14/21 samples and the only strain in 12. Complete genomes were determined for all strains, including isolates of persistent strains from their first and last appearances. Persistent strains belonged to phylogroups B2 (ST1236 and ST420), A (ST10 and ST93), or D (ST362). Single-sample strains were phylogroup A, B1, B2, D or F. Confirming extended survival in the gut, only minor changes were detected in the chromosomes of persistent strains over their residence periods. However, one lost a plasmid and two gained plasmids, providing evidence for horizontal gene transfer in the gut. A Z-type multidrug resistance plasmid moved from one persistent strain to the next and exchanged a small backbone region with a plasmid in the recipient. An M-type arsenic resistance plasmid was acquired by two different strains and transferred to laboratory *E. coli* with two co-resident plasmids. All persistent strains, but only 2/10 single-sample strains, carried one or more plasmid-borne bacteriocin determinants.

## Introduction

*Escherichia coli*, a facultative anaerobe that primarily resides in the intestinal tracts of vertebrates, is carried by most humans throughout life.^1,2^ However, *E. coli* can also cause disease at intestinal and extraintestinal sites.^3,4^ It is the leading cause of community-acquired urinary tract infections (UTIs) globally and also causes bacteraemia and neonatal meningitis.^3–5^ Treatment of these infections can be complicated by extensive antibiotic resistance, and *E. coli* is a leading cause of resistance-associated deaths.^6^

To date, *E. coli* has received far more attention as a causative agent of human disease than it has in its primary niche as a gastrointestinal resident. *E. coli* is a highly diverse species, and studies of human commensal *E. coli*, here designated ComEC, over the past century have been limited in their ability to resolve *E. coli* strains by the typing methods available at the time. This issue was recently reviewed by Martinson and Walk.^2^ Currently, *E. coli* is usually divided in order of increasing resolution into phylogenetic groups, sequence types, and strains, and it is clear that all phylogroups are well represented in ComEC populations. A 2010 compilation of 1,889 ComEC isolates derived from 25 studies on multiple continents found that, though their relative abundances varied across studies, overall 32.0% were phylogroup A, 29.4% B2, 20.7% D, and 17.9% B1.^7^ More recent studies of ComEC isolated in France and Australia also found that all phylogroups were represented,^8,9^ including groups C and F that were defined in 2013.^10^ ComEC also represents a reservoir for antibiotic resistance genes,^11^ and although the human gut has long been considered a site where horizontal gene transfer can occur,^12–14^ historic studies of transfer in the gut have been limited in their ability to examine the plasmids responsible and distinguish bacterial hosts involved.

Most studies of ComEC involve single faecal samples or rectal swabs and look at only a single colony per sample,^8,15^ providing no strain abundance information. Consequently, only a few studies have examined how many different *E. coli* are present simultaneously, finding 1-6 different strains per sample.^9,11,16–18^ Considerably less is known about the lengths of time specific *E. coli* strains reside in the healthy human gut. Tracking strain residency lengths and dynamics is challenging, requiring longitudinal sampling of human subjects and the ability to detect and accurately resolve *E. coli* strains within and between samples. Historic studies of ComEC residency using a variety of methodologies have been reviewed by Martinson and Walk, who surmised that: individual humans carry many *E. coli* strains over their lifetime, new strains are ingested constantly, and most are present only transiently before being lost at the rate of gut transit.^2^ Other strains establish themselves and can persist in the gut for various lengths of time, with estimates ranging from days to weeks, months, or years.^2^ Hence, gut residency is clearly a dynamic process. There is still very limited understanding of the factors that shape *E. coli* gut residency dynamics, which might include competition for space and nutrients, host adaptation, or the capacity to kill competing strains.^2^ The ability to investigate these traits has been constrained by the paucity of ComEC strain collections, and particularly ones tied to rigorously verified longitudinal residency data.

Few molecular era studies have examined the dynamics of ComEC populations carried by healthy adults.^17–23^ The most comprehensive tracked Enterobacteriaceae carried by eight healthy adults in the United States, with faecal samples collected at regular two-week intervals over periods of 245 to 849 days and up to 95 isolates per sample examined.^17^ This yielded a total of 37 resident strains, 34 of which were *E. coli*, exhibiting highly variable residency periods that ranged from tens to hundreds of days.^17^ Patterns of residency and turnover differed markedly between subjects,^17^ and similar outcomes were seen in a cohort of 11 adults over a 4-year period,^19^ and a cohort of 46 adults over a 5-month period.^21^ In another study the *E. coli* population carried by a single healthy adult was examined over a 20-year period, with 22 faecal samples collected at highly irregular intervals and 10 isolates examined per sample.^18,24^ A total of five ‘resident’ strains with residency periods of 7, 28, 315, 489 and 1,065 days were recovered and their complete genome sequences have been determined.^18^

Whole genome sequencing is the latest available tool for the characterisation of ComEC. Long-read sequencing data facilitates the assembly of complete genome sequences that provide information on the structures of chromosomes and plasmids.^25–27^ Complete genomes should enable detailed investigation of the genomic traits associated with successful gut residency, and precise tracking allowing for definitive characterisation of horizontal gene transfer in the gut. However, there are currently very few complete genome sequences for ComEC strains associated with gut residency data. Some studies have generated complete genome sequences for ComEC strains,^15,28–31^ but as far as we are aware, the five resident strains mentioned above are the only examples of complete genomes tied to residency data. There is therefore a clear need to generate high-quality complete genome assemblies for more verified long-term resident ComEC strains.

We have previously developed a detailed protocol for the isolation and characterisation of ComEC that was specifically designed to detect the range of strains present and hence facilitate the study of longitudinal strain dynamics and detection of horizontal gene transfer events occurring in the gut.^11^ The protocol involves examining 100 *E. coli* isolates per faecal sample using phenotypic and molecular methods to distinguish strains (Figure 1). Here, we report the composition and ecological dynamics of the ComEC population present in one individual using this protocol. Subject 24 provided multiple samples at regular intervals over a 700-day period between 2012 and 2014, and did not take antibiotics or visit a hospital throughout sampling. As there is currently no agreed definition of the length of residency that constitutes persistence in the gut, we classified strains that were present in consecutive samples (minimum 20 days apart) as ‘persistent’. Complete genomes were determined for all Subject 24 *E. coli* strains. In order to unambiguously confirm persistence and determine if differences had accrued over the courses of their residencies, an isolate from the first and last appearance of each persistent strain was sequenced and compared. Plasmid transfer and acquisition events in the healthy human gut were characterised.

**Figure 1:**
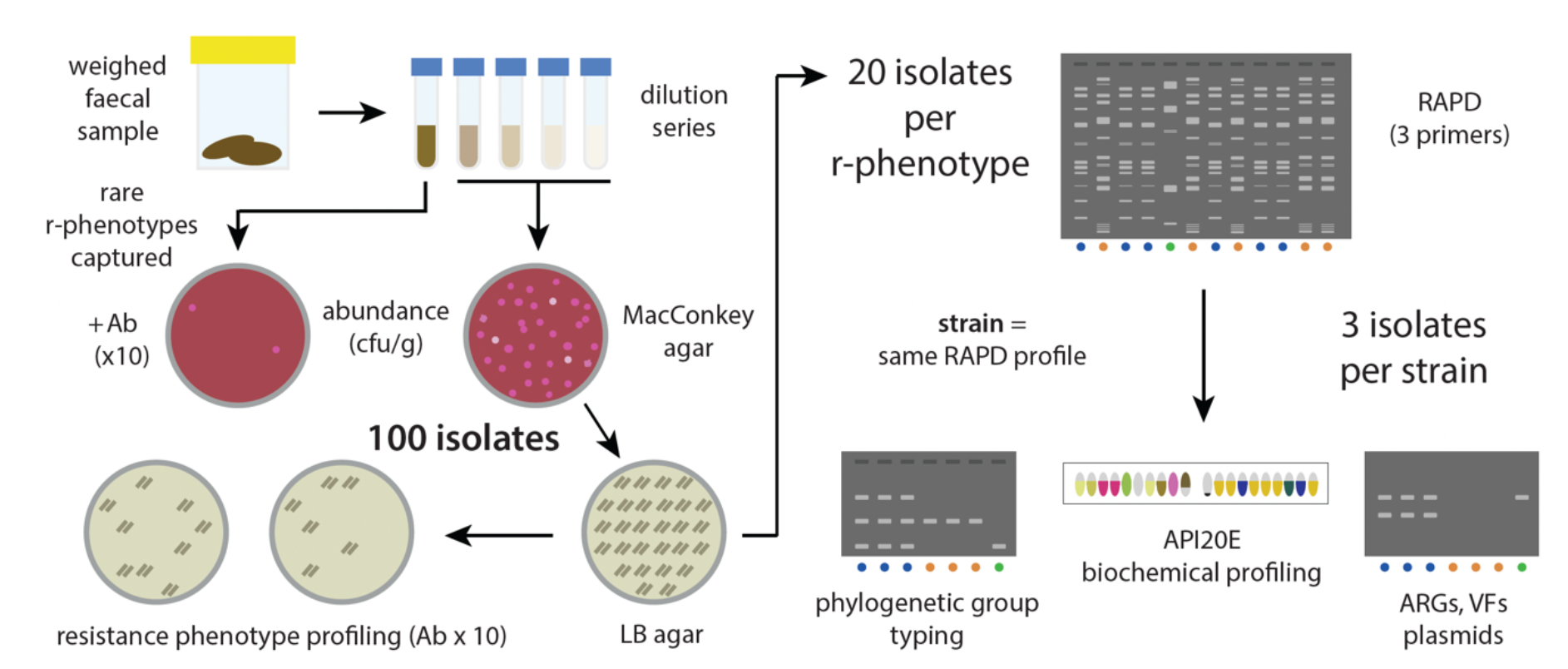
Protocol for isolation and characterisation of commensal *E. coli*. Overview of the steps taken to isolate *E. coli* from faecal samples, identify strains, and characterise representative isolates by molecular and biochemical methods. Abbreviations used: Ab = antibiotic; r-phenotype = resistance phenotype; RAPD = random amplification of polymorphic DNA; cfu/g = colony forming units per gram.

## Methods

### Sampling rationale and strategy

This study was conducted with informed consent and approved by the University of Sydney Human Research Ethics Committee (04-2008/10778). Fresh faecal samples were collected at approximately monthly intervals over a 700-day period.^32^

### Isolation and characterisation of *E. coli*

*E. coli* was isolated and characterised as described previously.^11^ Briefly, freshly voided (≤15 minutes) faecal samples were weighed and 100 µL volumes of 10^-3^ to 10^-7^ dilutions in sterile 0.9% saline were spread in duplicate on MacConkey agar (Oxoid) and incubated at 37°C overnight. Colony counts from MacConkey agar were used to determine the abundance (colony forming units per gram; cfu/g) of Enterobacteriaceae in each sample. To detect rare resistance phenotypes, 10^-2^ dilutions were plated on MacConkey agar containing ampicillin (100 µg/mL), cefotaxime (1 µg/mL), ceftazidime (1 µg/mL), chloramphenicol (25 µg/mL), kanamycin (20 µg/mL), spectinomycin (25 µg/mL), streptomycin (1 µg/mL), or tetracycline (10 µg/mL), and Mueller-Hinton agar (Oxoid) containing sulphamethoxazole (100 µg/mL), or trimethoprim (25 µg/mL).

100 presumptive *E. coli* colonies were picked at random from antibiotic-free MacConkey agar plates and patched across the series of antibiotic-containing plates listed above to assign resistance phenotypes. Isolates that grew on Simmons Citrate Agar were considered unlikely to be *E. coli* and characterised separately.

Metabolic typing and various PCR-based methods were used to differentiate and characterise *E. coli* strains as described previously.^11^ Briefly, three isolates of each antibiotic resistance phenotype, and 20 antibiotic-susceptible isolates from each sample were subjected to random amplification of polymorphic DNA (RAPD) using three different primers. Isolates of the same phenotype that produced identical RAPD patterns with all three primers were considered the same strain. Three representatives of each strain from each sample were biochemically profiled using API20E strips (bioMérieux), confirmed to be *E. coli* by PCR with primers targeting the *uidA* gene,^33^ assigned to phylogroups using a multiplex PCR,^34^ and screened for plasmid replicons as described previously.^35^

### Plasmid transfer and molecular characterisation

Plasmid transfer ability was assessed in mating experiments performed on agar plates as described previously using ComEC strains as donors and the nalidixic acid-resistant *E. coli* UB5201 as a recipient.^35^ Subsequent UB5201 transconjugants were mated with streptomycin-resistant or rifampicin-resistant *E. coli* UB1637 to confirm independent transfer ability and assess transfer frequencies. Transconjugants were selected on media containing 25 µg/mL nalidixic acid for UB5201, 1 µg/mL streptomycin or 100 µg/mL rifampicin for UB1637, and the relevant antibiotic (concentrations listed above) or 30 mM sodium arsenate for the plasmid of interest.

Plasmid DNA was isolated by alkaline lysis and visualised on agarose gels after pulsed-field gel electrophoresis by post-staining with ethidium bromide (5 µg/mL) as described previously using plasmid DNA from *E. coli* 39R861+ as a size standard.^36^ Restriction digests were performed and visualised as described previously.^37^

### Genome sequencing, assembly and analysis

Genomic DNA was extracted using the Qiagen UltraClean kit according to the manufacturer’s instructions and sequenced on the Illumina and Oxford Nanopore Technologies platforms at Plasmidsaurus. Illumina sequence data for five antibiotic resistant strains was obtained previously.^38^ Combined short and long reads were assembled using hybracter.^39^ Phylogroup typing was performed using ezclermont (github.com/nickp60/ezclermont). Antibiotic resistance genes were detected using ResFinder.^40^ Plasmid replicons were detected using PlasmidFinder,^41^ with small plasmids categorised according to their mode of replication: theta-replicating RNA-initiating (θ-RNA), theta-replicating Rep protein-initiating (θ-Rep) or rolling-circle (RC), as previously.^42,43^ FII and FIB-type replicons were further sub-typed using the replicon allele database hosted at PubMLST (pubmlst.org/organisms/plasmid-mlst), and M and Z-type replicons as described previously.^35,44^ Insertion sequences were identified using the ISFinder database.^45^ Sequences were compared to one another and to sequences in GenBank using BLAST.^46^

### Data availability

Sequencing reads and complete genome assemblies for all strains, including 24.21-Cfr and 24.21-Kva, are available from NCBI under BioProject accession PRJNA1474251.

## Results

### Overview of Subject 24 *E. coli* carriage

Faecal samples were collected from a single healthy adult, Subject 24, who had not taken any antibiotics for at least three years prior to sampling and did not take antibiotics throughout the sampling period. Twenty-one samples were collected at approximately monthly intervals over a 700-day period between 2012 and 2014, save for a 3-month gap between samples on days 171 and 282 (Figure 2). *E. coli* was isolated from all samples, and direct plating revealed its abundance was largely consistent across samples, ranging from 7.3x10^6^ to 5.7x10^7^ cfu/g (Table S1). This falls within the wider range of 2.0x10^4^ to 1.7x10^8^ cfu/g reported for a cohort of 20 healthy adults sampled by the same method.^11^ In general, *E. coli* was the only species of Enterobacteriaceae detected. However, one strain each of *Citrobacter freundii* (24.21-Cfr) and *Klebsiella variicola* (24.21-Kva) were isolated from sample 21 (Table S1).

**Figure 2:**
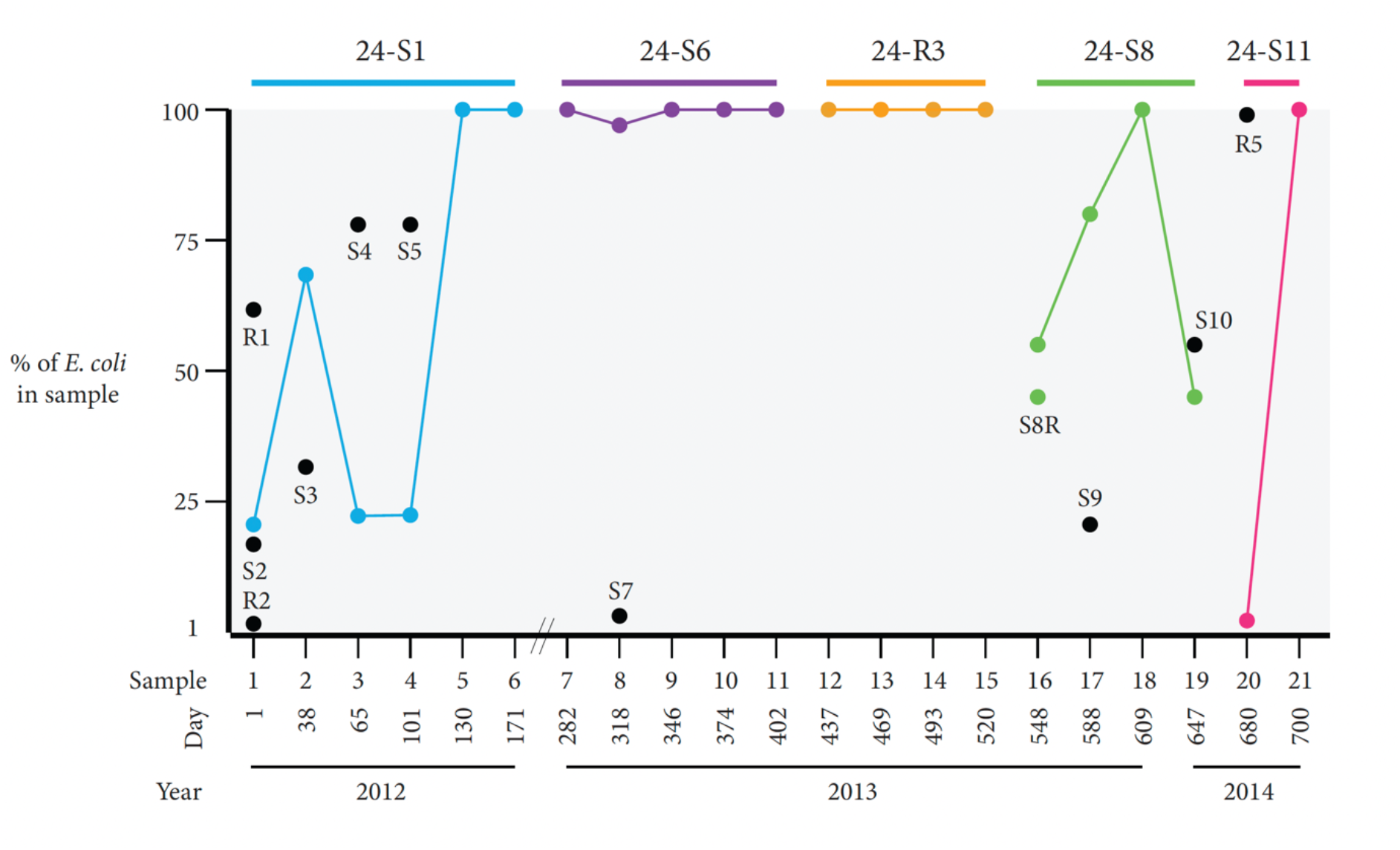
Relative abundance of commensal *E. coli* strains in Subject 24. Graphed abundances of *E. coli* strains isolated from each of 21 faecal samples from Subject 24, shown as a percentage of *E. coli* detected per sample. Coloured dots linked by coloured lines represent persistent strains with their identities labelled above, while black dots represent single-sample strains with their identities indicated using the last part of their strain names (e.g., 24.1-R1 is labelled “R1”).

To examine strain diversity, 100 presumptive *E. coli* colonies per sample were screened using the workflow shown in Figure 1 to identify closely related isolates. Isolates were classified initially by antibiotic resistance phenotype, then as belonging to the same strain if a subset of isolates representing each resistance type (three for each, 20 for antibiotic susceptible) exhibited the same RAPD profiles. Three isolates representing each putative strain in each sample were characterised further, with shared phylogroup and API20E profiles supporting strain identities. The number of strains detected this way was one or two in all but the first sample, which included three. An additional rare antibiotic-resistant strain was identified in sample 1 by direct plating on antibiotic-containing media (Figure 1), and its abundance was estimated at 0.9% of *E. coli* in that sample (Table S1). Within samples, individual strains accounted for 1-100% of *E. coli* (Table S1), and any strain that accounted for >80% of the *E. coli* in a sample was considered ‘dominant’.

In total, only 15 unique strains were recovered and five were antibiotic-resistant. Five strains, hereafter referred to as ‘persistent’, were present in multiple samples. Of the 10 found in a single sample each, only seven can be considered potentially ‘transient’ (resided <2 months) as the first sample yielded three strains for which prior extended residence cannot be ruled out. Residence times for persistent strains ranged from ≥83 to ≥170 days, but only ≥20 days for the strain that was present in the final two samples.

### Characteristics of persistent *E. coli* strains

The first persistent strain, 24-S1 (phylogroup B2), was recovered from six faecal samples over a six-month period in 2012, equating to a residency of ≥170 days. The relative abundance of 24-S1 varied across this period, accounting for <25% of *E. coli* in samples 1, 3 and 4, 70% in sample 2, and all detected *E. coli* in samples 5 and 6 (Figure 2). 24-S1 disappeared in the approximately three-month gap between samples 6 (day 171) and 7 (day 282).

The next two persistent strains, 24-S6 (phylogroup B2) and 24-R3 (phylogroup A) were consistently dominant, accounting for almost all *E. coli* in the samples they appeared in (Figure 2). 24-S6 replaced 24-S1 in the 111 days between samples 6 and 7, and was the only strain found in sample 7. It persisted for at least four months (≥120 days; <266 days) until sample 11 (day 402), and only a single additional strain was isolated during its residency, namely 24.8-S7 which accounted for 5% of *E. coli* in sample 8 (Figure 2). 24-R3, which was resistant to ampicillin, streptomycin, sulphamethoxazole and trimethoprim (ApSmSuTp) replaced 24-S6 in the 35-day interval between samples 11 and 12 and persisted for at least three months (≥83 days; <146 days). It was the only strain isolated from samples 12 to 15 (Figure 2).

In sample 16 (day 548), 55/100 isolates were resistant to ApSmSuTp, the same phenotype as previously persistent strain 24-R3. However, the ApSmSuTp resistant isolates in sample 16 that were tested further (n=3) exhibited RAPD patterns distinct from those of 24-R3 and hence represented a new strain. As not all ApSmSuTp resistant isolates were tested further, we cannot exclude the possibility that 24-R3 was present in this sample. The remaining 45 isolates in sample 16 were antibiotic sensitive but all of those tested (n=20) had the same RAPD pattern as the ApSmSuTp resistant isolates. Thus, sample 16 contained two phenotypic variants of the same strain that were designated 24-S8 and 24-S8R. The ApSmSuTp resistance phenotype was not detected again after sample 16, but 24-S8 (phylogroup A) was resident for ≥99 days through to sample 19 (day 647). While 24-S8 was the only strain in samples 16 (with 24-S8R) and 18, and was the dominant strain in sample 17 (80%), it accounted for just 45% of *E. coli* in sample 19 and was not seen 33 days later in sample 20 on day 680 (Figure 2).

The final persistent strain, 24-S11 (phylogroup D), was only isolated from samples 20 and 21 at the end of the sampling period. Detection of 24-S11 in sample 20 was fortuitous, as just one colony among the 100 screened was sensitive to tetracycline (Table S1). Over the next 19 days, 24-S11 rose in abundance to account for all *E. coli* tested from sample 21 (day 700). The ≥20-day observed residency period of 24-S11 was significantly shorter than those of the four previous persistent strains as sample 21 was the final sample. Hence, we cannot exclude ongoing residence of 24-S11 beyond the end of the study.

### Characteristics of persistent strain genomes

To characterise the five persistent strains and verify their persistence, complete genome sequences were determined for an isolate from both the first and last sample in which they were detected. An isolate representing 24-S8R, the antibiotic-resistant variant of 24-S8, was also sequenced. Features of the 11 complete genomes for the persistent strains are outlined in Table 1. Each genome contained a chromosome (4.71 - 4.89 Mbp) and 2 to 6 plasmids. Each persistent strain was a different sequence type, namely ST1236 (24-S1), ST420 (24-S6), ST10 (24-R3), ST93 (24-S8) and ST362 (24-S11).

**Table 1:** Genomic characteristics of Subject 24 persistent *E. coli* strains.

| Strain | Sample | Phylo Group | ST | DNA molecule |  | NCBI BioSample # |
|---|---|---|---|---|---|---|
|  |  |  |  | Size (bp) | Type |  |
| 24-S1 | 1 | B2 | 1236 | 4,792,181 | chromosome | SAMN60589304 |
|  |  |  |  | 140,874 | FII:FIB plasmid |  |
|  |  |  |  | 93,621 | phage-plasmid |  |
|  |  |  |  | 33,809 | X4 plasmid |  |
| 24-S1 | 6 | B2 | 1236 | 4,792,176 | chromosome | SAMN60589305 |
|  |  |  |  | 140,874 | FII:FIB plasmid |  |
|  |  |  |  | 33,809 | X4 plasmid |  |
| 24-S6 | 7 | B2 | 420 | 4,738,296 | chromosome | SAMN60589312 |
|  |  |  |  | 194,715 | FII:FIB plasmid |  |
|  |  |  |  | 100,871 | phage-plasmid |  |
|  |  |  |  | 45,425 | phage-plasmid |  |
|  |  |  |  | 5,729 | θ-Rep plasmid |  |
|  |  |  |  | 2,311 | RC plasmid |  |
| 24-S6 | 11 | B2 | 420 | 4,738,284 | chromosome | SAMN60589313 |
|  |  |  |  | 194,715 | FII:FIB plasmid |  |
|  |  |  |  | 100,871 | phage-plasmid |  |
|  |  |  |  | 45,425 | phage-plasmid |  |
|  |  |  |  | 5,729 | θ-Rep plasmid |  |
|  |  |  |  | 2,311 | RC plasmid |  |
| 24-R3 | 12 | A | 10 | 4,970,131 | chromosome | SAMN60589315 |
|  |  |  |  | 104,029 | Z plasmid |  |
|  |  |  |  | 76,379 | FIB plasmid |  |
|  |  |  |  | 6,646 | θ-RNA plasmid |  |
|  |  |  |  | 6,217 | θ-Rep plasmid |  |
|  |  |  |  | 5,165 | θ-Rep plasmid |  |
|  |  |  |  | 4,072 | θ-Rep plasmid |  |
| 24-R3 | 15 | A | 10 | 4,970,131 | chromosome | SAMN60589316 |
|  |  |  |  | 104,029 | Z plasmid |  |
|  |  |  |  | 76,379 | FIB plasmid |  |
|  |  |  |  | 6,646 | θ-RNA plasmid |  |
|  |  |  |  | 6,217 | θ-Rep plasmid |  |
|  |  |  |  | 5,165 | θ-Rep plasmid |  |
|  |  |  |  | 4,072 | θ-Rep plasmid |  |
| 24-S8 | 16 | A | 93 | 4,708,120 | chromosome | SAMN60589317 |
|  |  |  |  | 68,957 | FII plasmid |  |
|  |  |  |  | 5,145 | θ-Rep plasmid |  |
| 24-S8R | 16 | A | 93 | 4,708,122 | chromosome | SAMN60589318 |
|  |  |  |  | 104,024 | Z plasmid |  |
|  |  |  |  | 68,957 | FII plasmid |  |
|  |  |  |  | 5,145 | θ-Rep plasmid |  |
| 24-S8 | 19 | A | 93 | 4,708,120 | chromosome | SAMN60589319 |
|  |  |  |  | 72,932 | M plasmid |  |
|  |  |  |  | 68,957 | FII plasmid |  |
|  |  |  |  | 5,145 | θ-Rep plasmid |  |
| 24-S11 | 20 | D | 362 | 4,886,305 | chromosome | SAMN60589322 |
|  |  |  |  | 68,957 | FII plasmid |  |
|  |  |  |  | 5,150 | θ-Rep plasmid |  |
| 24-S11 | 21 | D | 362 | 4,886,308 | chromosome | SAMN60589323 |
|  |  |  |  | 72,932 | M plasmid |  |
|  |  |  |  | 68,957 | FII plasmid |  |
|  |  |  |  | 5,150 | θ-Rep plasmid |  |

In all cases, the chromosomes of pairs of isolates representing the same persistent strain exhibited ≤6 single-nucleotide differences or small indels (detailed in Table S2). For example, the 24-S1 chromosome exhibited six differences between samples 1 and 6: five single-base changes and a 4 bp deletion in the *wcaA* glycosyltransferase gene that features 35 copies of the same 4 bp motif. Persistent strain chromosomes were structurally identical at the beginning and end of their residencies, with no large deletions, inversions or differences in the numbers or positions of insertion sequences. This near-perfect chromosomal conservation unambiguously confirmed persistence of these five strains in the gut of Subject 24.

The plasmid content of persistent strains was more dynamic. Three of the five strains exhibited a change in plasmid content over the course of their residencies. 24-S1 isolated from sample 1 carried the 93,621 bp phage-plasmid pCESC7, which was not present in the 24-S1 genome from sample 6. This was the only case of plasmid loss. The remaining events involved the acquisition of plasmids, consistent with plasmid transfer in the gut of Subject 24.

### Subject 24 *E. coli* hosted a diverse array of plasmids

Complete genome sequences were determined for the strains isolated from only a single sample each. These also exhibited diverse phylogroups, sequence types, and plasmid content (Table 2). Two single-sample strains, 24.1-R1 (phylogroup F; ST1485) and 24.2-S3 (phylogroup D; ST38) did not contain plasmids. Other strains carried one to six plasmids each (Tables 1; 2).

**Table 2:**
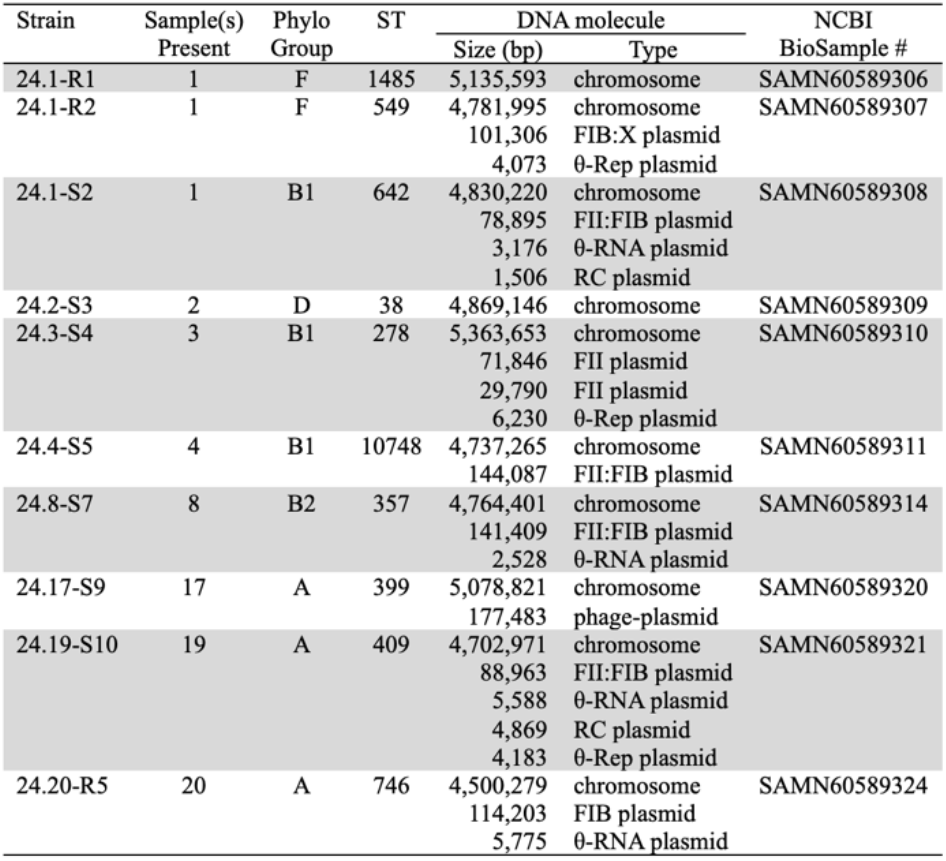
Genomic characteristics of Subject 24 single-sample *E. coli* strains.

| Strain | Sample(s) Present | Phylo Group | ST | DNA molecule |  | NCBI BioSample # |
|---|---|---|---|---|---|---|
|  |  |  |  | Size (bp) | Type |  |
| 24.1-R1 | 1 | F | 1485 | 5,135,593 | chromosome | SAMN60589306 |
| 24.1-R2 | 1 | F | 549 | 4,781,995 | chromosome | SAMN60589307 |
|  |  |  |  | 101,306 | FIB:X plasmid |  |
|  |  |  |  | 4,073 | θ-Rep plasmid |  |
| 24.1-S2 | 1 | B1 | 642 | 4,830,220 | chromosome | SAMN60589308 |
|  |  |  |  | 78,895 | FII:FIB plasmid |  |
|  |  |  |  | 3,176 | θ-RNA plasmid |  |
|  |  |  |  | 1,506 | RC plasmid |  |
| 24.2-S3 | 2 | D | 38 | 4,869,146 | chromosome | SAMN60589309 |
| 24.3-S4 | 3 | B1 | 278 | 5,363,653 | chromosome | SAMN60589310 |
|  |  |  |  | 71,846 | FII plasmid |  |
|  |  |  |  | 29,790 | FII plasmid |  |
|  |  |  |  | 6,230 | θ-Rep plasmid |  |
| 24.4-S5 | 4 | B1 | 10748 | 4,737,265 | chromosome | SAMN60589311 |
|  |  |  |  | 144,087 | FII:FIB plasmid |  |
| 24.8-S7 | 8 | B2 | 357 | 4,764,401 | chromosome | SAMN60589314 |
|  |  |  |  | 141,409 | FII:FIB plasmid |  |
|  |  |  |  | 2,528 | θ-RNA plasmid |  |
| 24.17-S9 | 17 | A | 399 | 5,078,821 | chromosome | SAMN60589320 |
|  |  |  |  | 177,483 | phage-plasmid |  |
| 24.19-S10 | 19 | A | 409 | 4,702,971 | chromosome | SAMN60589321 |
|  |  |  |  | 88,963 | FII:FIB plasmid |  |
|  |  |  |  | 5,588 | θ-RNA plasmid |  |
|  |  |  |  | 4,869 | RC plasmid |  |
|  |  |  |  | 4,183 | θ-Rep plasmid |  |
| 24.20-R5 | 20 | A | 746 | 4,500,279 | chromosome | SAMN60589324 |
|  |  |  |  | 114,203 | FIB plasmid |  |
|  |  |  |  | 5,775 | θ-RNA plasmid |  |

In total, the 15 strains hosted a set of 34 plasmids that ranged in size from 1,506 bp to 194,715 bp and were of diverse replicon types (Table 3). Almost half (n=16) were small plasmids <10 kbp. Amongst the large plasmids the most common were F-types (FII or FIB replicons), which were found in 12/15 strains and accounted for 11/34 plasmids in the set. The next most common replicon types were amongst the small plasmids, θ-Rep (8/34) and θ-RNA (5/34).

**Table 3:** Plasmids in Subject 24 *E. coli*.

| Replicon type |  | Plasmid | Size (bp) | Host(s) | Col/Mcc | Resistance conferred<sup>7</sup> |
|---|---|---|---|---|---|---|
| Broad | Specific |  |  |  |  |  |
| F | FII-64:FIB-62<sup>1</sup> | pCESC3 | 140,874 | 24-S1; 24.8-S7<sup>4</sup> | V, Ia, J25 | - |
|  | FIB-54<sup>1</sup> | pCESC9 | 78,895 | 24.1-S2 | - | - |
|  | FII-63 | pCESC12 | 71,846 | 24.3-S4 | - | - |
|  | FII<sup>3</sup> | pCESC13 | 29,790 | 24.3-S4 | - | - |
|  | FII-56:FIB-62<sup>2</sup> | pCESC15 | 144,087 | 24.4-S5 | - | - |
|  | FII-18:FIB-1 | pCESC16 | 194,715 | 24-S6 | V | - |
|  | FIB-64 | pCESC22 | 76,378 | 24-R3 | - | - |
|  | FII-4 | pCESC6 | 68,957 | 24-S8; 24-S11 | B17 | - |
|  | FII-100:FIB(K) | pCESC28 | 88,963 | 24.19-S10 | - | - |
| Cointegrate | FIB(K):X1 | pCERC17 | 101,306 | 24.1-R2 | - | ApCmSmSpSuTcTp |
|  | FIB:Y | pCERC18 | 114,203 | 24.20-R5 | - | Tc |
| M | M1a | pCESC2 | 72,932 | 24-S8M; 24-S11M | - | arsenic |
| X | X4 | pCESC4 | 33,809 | 24-S1 | - | - |
| Z | Z-2 | pCERC12 | 104,029 | 24-R3; 24-S8R<sup>5</sup> | - | ApSmSuTp |
| phage-plasmid | p0111 | pCESC7 | 93,621 | 24-S1 | - | - |
|  | Y | pCESC17 | 100,871 | 24-S6 | - | - |
|  | - | pCESC18 | 45,425 | 24-S6 | - | - |
|  | Y | pCESC27 | 177,483 | 24.17-S9 | - | - |
| θ-RNA | - | pCESC8 | 4,073 | 24.1-R2 | - | - |
|  | - | pCESC10 | 3,176 | 24.1-S2 | - | - |
|  | - | pCESC21 | 2,528 | 24.8-S7 | - | - |
|  | - | pCESC23 | 6,646 | 24-R3 | E1 | - |
|  | - | pCESC29 | 5,588 | 24.19-S10 | - | - |
|  | - | pCESC32 | 5,775 | 24.20-R5 | - | - |
| θ-Rep | - | pCESC14 | 6,230 | 24.3-S4 | E7 | - |
|  | - | pCESC19 | 5,729 | 24-S6 | - | - |
|  | - | pCESC24 | 6,217 | 24-R3 | - | - |
|  | - | pCESC25 | 5,165 | 24-R3 | - | - |
|  | - | pCESC26 | 4,072 | 24-R3 | - | - |
|  | - | pCESC5 | 5,145 | 24-S8; 24-S11<sup>6</sup> | - | - |
|  | - | pCESC31 | 4,183 | 24.19-S10 | - | - |
| rolling-circle | - | pCESC11 | 1,506 | 24.1-S2 | - | - |
|  | - | pCESC20 | 2,311 | 24-S6 | - | - |
|  | - | pCESC30 | 4,869 | 24.19-S10 | - | - |

<sup>1</sup> One nucleotide difference relative to the reference sequence in PubMLST.
<sup>2</sup> Three nucleotide differences relative to the reference sequence in PubMLST.
<sup>3</sup> No matches to the PubMLST database.
<sup>4</sup> pCESC3-v2 variant in 24.8-S7 is 141,409 bp.
<sup>5</sup> pCERC12-v2 variant in 24-S8R is 104,024 bp.
<sup>6</sup> pCERC5-v2 variant in 24-S11 is 5,150 bp.
<sup>7</sup> Antibiotic abbreviations: ampicillin (Ap), chloramphenicol (Cm), streptomycin (Sm), spectinomycin (Sp), sulphamethoxazole (Su), tetracycline (Tc), trimethoprim (Tp).

Cases where identical or near-identical plasmids were carried by different strains are noted in Table 3 and described further in the sections below.

### Transfer of a multidrug resistance plasmid during persistent strain turnover

Both 24-R3 and 24-S8R were previously shown to transfer the ApSmSuTp resistance phenotype along with the same Z-type plasmid replicon to laboratory *E. coli* UB5201, suggesting that transfer of this plasmid had occurred in the gut of Subject 24.^35^ Here, the Z-type plasmid derived from 24-S8R transferred from UB5201 to plasmid-free *E. coli* UB1637 at a frequency of 1.7 x 10^-4^ transconjugants per donor (TC/D), confirming its ability to transfer independently.

It is possible that 24-R3 and 24-S8 were present together at some point in the 27 days between samples 15 and 16 (Figure 2), and that transfer of this Z-type plasmid from 24-R3 to 24-S8 generated the resistant variant 24-S8R (Figure 3A). Complete genome sequences confirmed that all resistance genes in 24-R3 and 24-S8R were located in identical complex resistance regions in a 104 kbp Z-type plasmid (Figure 3B). Consistent with its transfer ability, the plasmid backbone contains complete and uninterrupted sets of conjugation determinants, including for the two conjugative pili that are characteristic of members of the I-complex.^47^ The sequences of this plasmid, pCERC12, from the first and last appearances of 24-R3 are identical. However, in 24-S8R the pCERC12 variant pCERC12-v2 is 5 bp smaller due to a patch in the backbone that is 129 bp in 24-R3 and 124 bp in 24-S8R (Figure 3B). This patch features 21 nucleotide differences including indels that account for the size difference. Further analysis revealed the 124 bp patch in pCERC12-v2 from 24-S8R is identical to part of the backbone of the FII-4 plasmid pCESC6 that was also present in 24-S8R. The patch is immediately flanked by 167 bp and 310 bp segments that are identical in pCESC6 and pCERC12, and these likely facilitated homologous recombination that introduced the 124 bp patch into pCERC12 after it was acquired by 24-S8. Hence, this difference is not inconsistent with *in situ* plasmid transfer.

**Figure 3:**
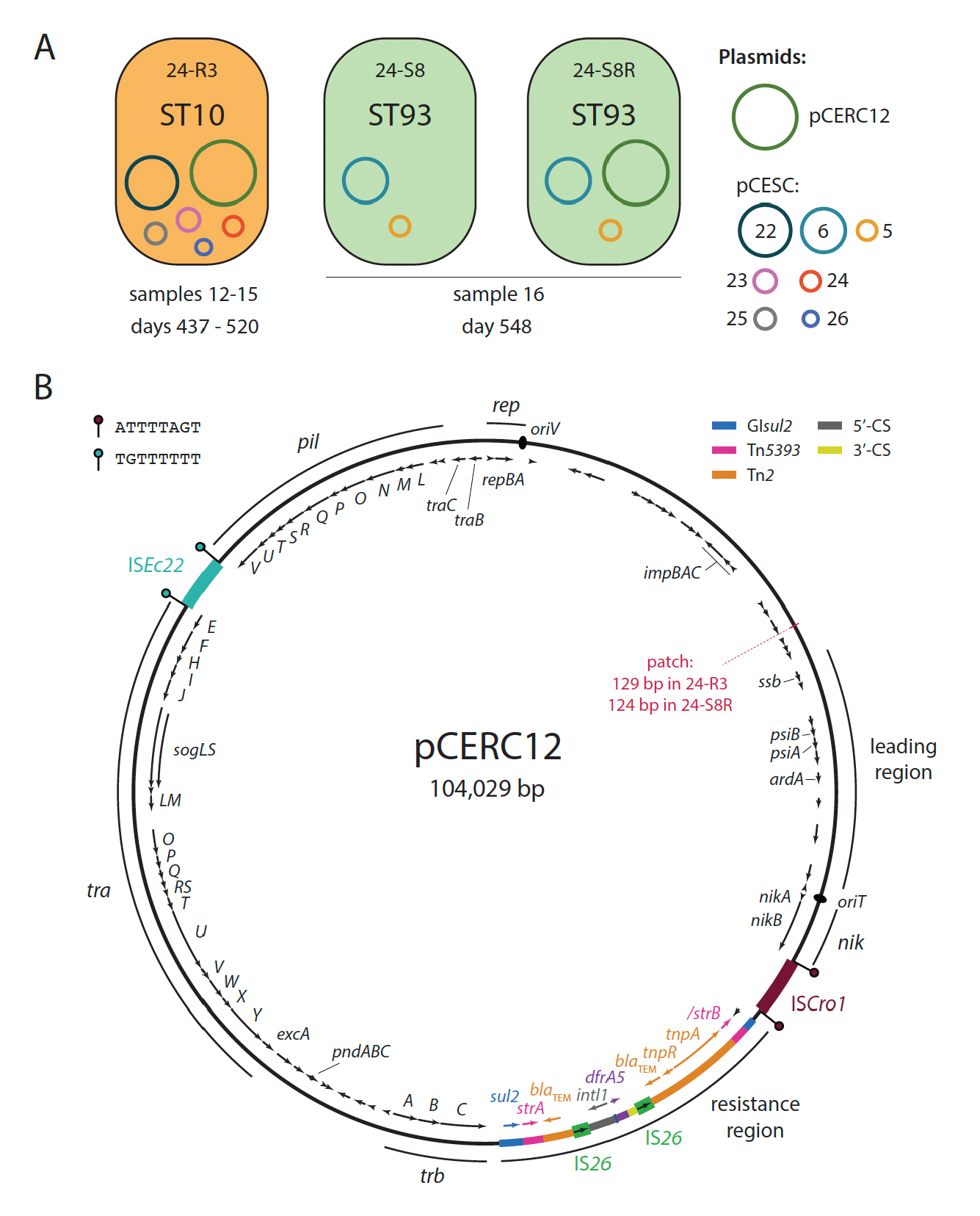
Transfer of Z-type plasmid pCERC12 in Subject 24. A) Plasmid content of 24-R3, 24-S8 and 24-S8R. Each is shown as an oval filled with the colour corresponding to that used to represent each strain in Figure 2, with isolation details below. Plasmids are shown as coloured circles within cells, with their identities indicated in the legend to the right. B) Circular map of pCERC12. Open reading frames are shown as labelled arrows inside the circle, and regions of the plasmid are marked by labelled arcs outside the circle. The positions of the origin-of-replication (*oriV*) and origin-of-transfer (*oriT*) are marked by labelled ovals. The identities of insertion sequences are indicated by labels outside the circle. The derivations of segments in the resistance region are denoted by colour as shown in the legend at top right. Target site duplications are shown as ‘lollipops’ flanking the element that generated them, with sequences indicated at top left. The location of the recombinant patch that distinguishes the pCERC12 variant found in 24-S8R is indicated by a labelled red line.

### Acquisition of an arsenic resistance plasmid by two persistent strains

Strain 24-S8 acquired another plasmid towards the end of its residency. Isolates of 24-S8 from sample 19 contained a 72,932 bp M-type plasmid, pCESC2, that was not present in 24-S8 or 24-S8R genomes from sample 16 and was not detected by PCR in 24-S8 isolates from samples 17 and 18 (Figure 4A). pCESC2 was present again in sample 21, where it was carried by 24-S11, but it was not present in the lone 24-S11 isolate from sample 20. Isolates of 24-S8 and 24-S11 from samples 19 and 21, respectively, that contained pCESC2 will for clarity be referred to as 24-S8M and 24-S11M. The source of pCESC2 in Subject 24 was not detected, but may have been present as a minority of the population that escaped isolation or characterisation by our protocol.

**Figure 4:**
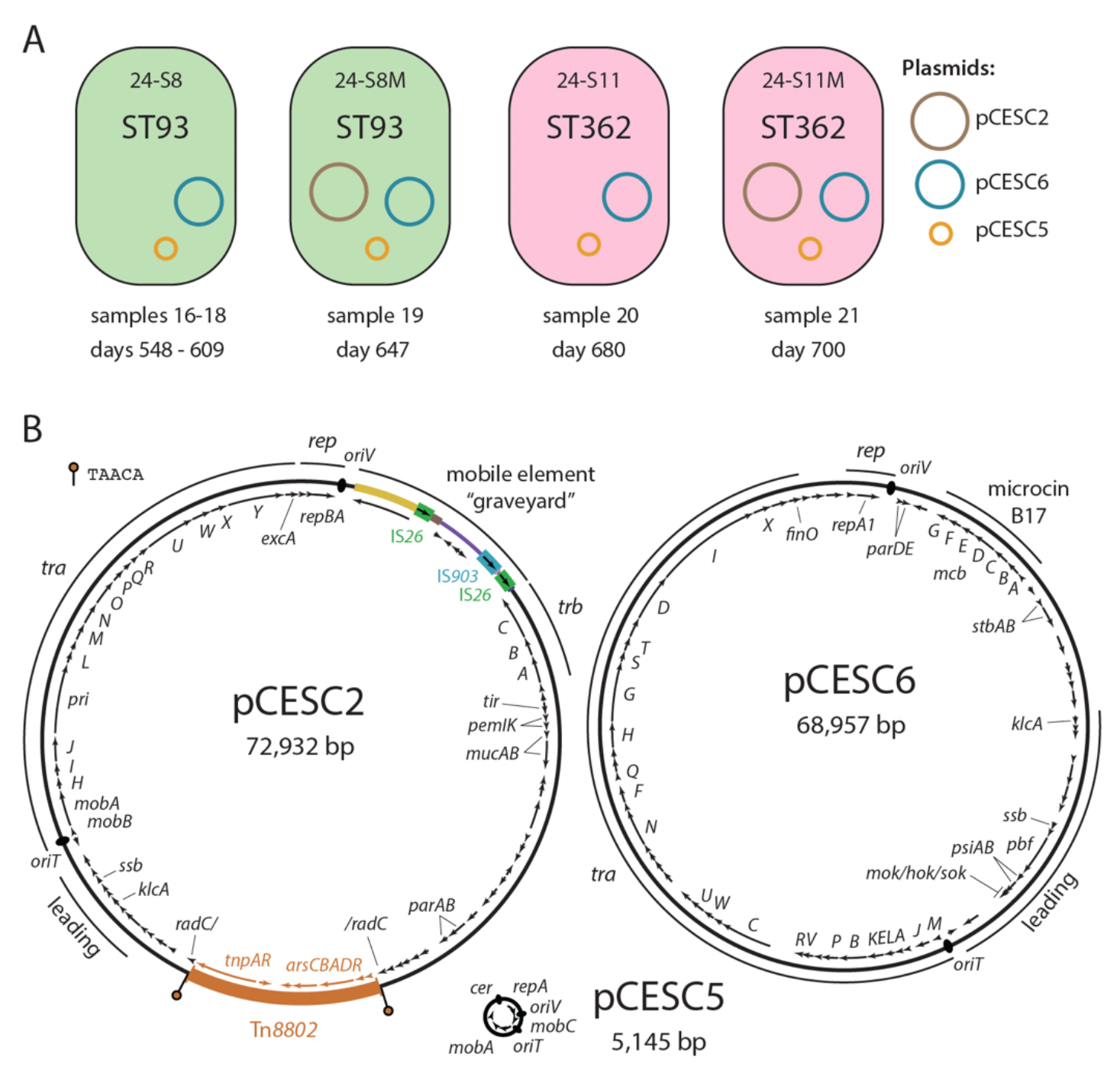
Acquisition of M-type plasmid pCESC2. A) Plasmid content of 24-S8, 24-S8M, 24-S11 and 24-S11M. Each is shown as an oval filled with the colour corresponding to that used to represent each strain in Figure 2, with isolation details below. Plasmids are shown as coloured circles within cells, with their identities shown in the legend to the right. B) Circular maps of pCESC2, pCESC6 and pCESC5, drawn to the same scale. Open reading frames are shown as labelled arrows inside the circle, and regions of the plasmid are marked by labelled arcs outside the circle. The positions of origin-of-replication (*oriV*), origin-of-transfer (*oriT*), and *cer* sites are marked by labelled ovals. Target site duplications are shown as ‘lollipops’ flanking the element that generated them, with sequences indicated at top left.

pCESC2 carries a complete transfer region and a complete copy of transposon Tn*8802*, a hybrid of TnARS and Tn*2* that has been reported to confer resistance to arsenite.^48^ The arsenic resistance phenotype was confirmed by comparing the growth of 24-S8/24-S11 and 24-S8M/24-S11M on media containing various concentrations of sodium arsenate or sodium arsenite. In both hosts, carriage of pCESC2 enabled growth on 80 mM sodium arsenate (20 mM without pCESC2) and 2.5 mM of sodium arsenite (<1 mM without pCESC2). To separate pCESC2 from co-resident plasmids, UB5201 was transformed with plasmid DNA extracted from 24-S11M and selected on 30 mM sodium arsenate. Restriction digests of plasmid DNA derived from UB5201 transformants confirmed that they only contained pCESC2. The transfer frequency of pCESC2 from UB5201 to plasmid-free UB1637 was very high, 6.7 x 10^-2^ TC/D.

### Co-transfer of pCESC2 with pCESC6 and pCESC5?

All isolates of 24-S8 and 24-S11 shared two further plasmids, the 69 kbp FII-4 pCESC6 and the 5.1 kbp θ-Rep pCESC5 (Figure 4A). pCESC6, which is identical in 24-S8 and 24-S11, is uninterrupted by MGEs and has a complete conjugative transfer region (Figure 4B). pCESC5 is a ColE2-like plasmid that was 5,145 bp in 24-S8 and 5,150 bp in 24-S11, with the size difference accounted for by single-base differences in the lengths of five different poly-T tracts. pCESC5 contains the replication initiation gene *repA*, a *cer*-like multimer resolution site, and an *oriT* site between *mobA* and *mobC* genes that together form a MOB_Q4_-type mobilisation system (Figure 4B).

When the conjugative ability of pCESC2 was tested in experiments using 24-S8M and 24-S11M as donors, rather than UB5201(pCESC2) as above, and plasmid-free UB5201 as recipient, each of the 18 arsenic-resistant UB5201 transconjugants examined further (three from each of three independent matings per donor strain) also contained pCESC6 and pCESC5. This suggests that these three plasmids may have transferred together from 24-S8M to a plasmid-free 24-S11 in the gut of Subject 24. However, the original host of pCESC2 in Subject 24 has not been identified, and we cannot rule out the possibility that 24-S8 and 24-S11 acquired pCESC2 from this unknown host independently. Either way, transfer of pCESC2 has clearly occurred in the gut.

### Plasmid-borne colicin determinants associated with persistence

We considered the possibility that bacteriocins may play an important role in the ability of *E. coli* strains to persist in the healthy human gut, owing to the competitive advantage they can confer.^49,50^ All five persistent strains, and only 2/10 single-sample strains, carried one or more plasmid-borne colicin or microcin determinants (Table 3).

Most bacteriocin genes were located in F-type plasmids. 24-S1 carried genes for colicin V (also called microcin V), colicin Ia and microcin J25 in the 140 kbp FII-64:FIB-62 plasmid pCESC3 (Figure S1). 24-S6 carried the colicin V genes in a different F-type plasmid, pCESC16 (FII-18:FIB-1). The 69 kbp FII-4 plasmid pCESC6 carried by 24-S8 and 24-S11 (Figure 4B) is colinear with and 97.6% identical to the well-studied microcin B17-producing plasmid pMccb17 (GenBank accession ON989342) that was originally found in a ComEC isolate from a healthy newborn in Spain in the 1970s.^51,52^ In the microcin B17-determining region, pCESC6 differs by just a single silent base change from pMccb17. The remaining persistent strain, 24-R3, carried genes for colicin E1 in the 6.6 kbp θ-RNA plasmid pCESC23, which is colinear with and 98.6% identical to the classic plasmid ColE1 (GenBank accession J01566) that was first reported in the 1960s,^53^ with most differences clustered in a 1.4 kbp recombinant patch containing the mobilisation determinants.

Transient strain 24.8-S7, the only phylogroup B2 strain in the single-sample group, carried determinants for colicin V, colicin Ia, and microcin J25 in a plasmid closely related to pCESC3. Relative to the 140,874 bp pCESC3 carried by 24-S1, the 141,409 bp pCESC3-v2 in 24.8-S7 had an additional 777 bp immediately adjacent to a copy of insertion sequence IS*1* and lacked a 258 bp backbone segment (Figure S1). The only other colicin plasmid found in a transient strain was pCESC14 (6,230 bp; θ-RNA) in 24.3-S4, which contains genes for colicin E7.

### Iron uptake systems

Siderophore systems in addition to the intrinsic enterobactin might provide a competitive advantage in gut colonisation and persistence.^23^ We therefore investigated the distribution of four acquired siderophore systems: aerobactin, salmochelin, yersiniabactin, and Sit. There was no clear association between siderophore gene carriage and extended gut residency (Table 4). Indeed, 24-S11 carried none. Rather, phylogroup B2 strains, whether persistent (24-S1 and 24-S6) or transient (24.8-S7), carried the most siderophore determinants. In all three B2 strains, siderophore determinants were found in the plasmids that contained the colicin determinants described above: pCESC16 in 24-S6 encoded salmochelin, aerobactin, and Sit, while pCESC3 in 24-S1 and pCESC3-v2 in 24.8-S7 encoded salmochelin. These B2 strains also carried chromosomal Sit genes, as well as genes for yersiniabactin in the chromosomal High Pathogenicity Island.^54^ Outside of phylogroup B2, all acquired siderophore determinants were chromosomal (Table 4).

**Table 4:** Siderophore determinants in Subject 24 *E. coli* strains.

| Strain | Phylo group | ST | Siderophore system<sup>1, 2</sup> |  |  |  |  |
|---|---|---|---|---|---|---|---|
|  |  |  | enterobactin | aerobactin | salmochelin | yersiniabactin | Sit |
| 24-S1 | B2 | 1236 | c | - | p | c | c |
| 24-S6 | B2 | 420 | c | p | p | c | p, c |
| 24-R3 | A | 10 | c | c | - | c | c |
| 24-S8 | A | 93 | c | - | - | - | c |
| 24-S11 | D | 362 | c | - | - | - | - |
| 24.1-S2 | B1 | 642 | c | - | - | - | - |
| 24.1-R1 | F | 1485 | c | - | - | - | c |
| 24.1-R2 | F | 549 | c | - | - | - | - |
| 24.2-S3 | D | 38 | c | - | - | - | c |
| 24.3-S4 | B1 | 278 | c | c | - | - | - |
| 24.4-S5 | B1 | 10748 | c | - | - | - | - |
| 24.8-S7 | B2 | 357 | c | - | p | c | c |
| 24.17-S9 | A | 399 | c | - | - | c | - |
| 24.19-S10 | A | 409 | c | - | - | - | c |
| 24.20-R5 | A | 746 | c | - | - | - | - |

<sup>1</sup> "c" denotes chromosomal location; "p" plasmid
<sup>2</sup> Genes detected for each system:
enterobactin [*entABCDEFS*, *fepABCDG*, *fes*], aerobactin [*iucABCD*, *iutA*], salmochelin [*iroBCDEN*], yersiniabactin [*ybtAEPQSTUX*, *fyuA*], Sit [*sitABCD*].

### Contexts of antibiotic resistance genes in Subject 24 strains

24.1-R1 was the only strain resistant to cephalosporins in our collection of 51 antibiotic-resistant ComEC strains isolated between 2008 and 2014.^38^ It did not carry any plasmids, and the two resistance determinants were found in MGEs inserted at different locations in the chromosome. The tetracycline resistance gene *tetA*(B) was in a complete copy of transposon Tn*10*, and the cephalosporin resistance gene *bla*_CMY-2_ was in a 3,709 bp IS*Ecp1* transposition unit that we described previously.^38^ All other antibiotic resistance genes in Subject 24 strains were located in plasmids, and the structure of these regions was clarified in the complete genomes. For example, 24.20-R5 carried the *tet*(A) tetracycline resistance gene in a 9,411 bp fragment of Tn*1721* that has been truncated by a copy of IS*26* in the 114,203 bp FIB-type plasmid pCERC18.

The resistance genes in Z plasmid pCERC12 were in a region derived from ones found in the same position in other I-complex plasmids, and made up of parts of the elements GI*sul2*, Tn*5393*, and Tn*2* that include the resistance genes *sul2*, *strAB* and *bla*_TEM_, respectively.^37,55,56^ In pCERC12, the region features a second copy of the *bla*_TEM_ gene and includes an IS*26*-bounded pseudo-compound transposon that contains a partial class 1 integron bearing the trimethoprim resistance gene *dfrA5* (Figure 3B).

24.1-R2 (phylogroup F; ST549) carried a 101,306 bp FIB(K):X1 cointegrate plasmid, pCERC17, that contained nine antibiotic resistance genes accounting for its phenotype. The resistance genes were located in a *sul3*-associated class 1 integron with cassette array *dfrA12*-orfF-*aadA2*-*cmlA1*-*aadA1*-*qacH*, and in segments derived from several other MGEs including Tn*2* (*bla*_TEM_), Tn*1721* (*tet*(A)), and Tn*5393* (*strAB*). pCERC17 also contained *mcr-9.1*, but this gene alone is not expected to confer resistance to colistin.^57^

## Discussion

Isolates from clinical sources account for the vast majority of *E. coli* genomes in public repositories,^58^ but the importance of also considering commensal strains in genomic evaluations of the species has been noted since the first complete ComEC genomes were published in 2008.^28,59^ Since that time, the enormous diversity amongst ComEC has been more widely appreciated,^8^ and it is clear that no single strain is representative of the ComEC population. Hence, a much larger collection of publicly available ComEC genomes, ideally complete genomes, will be essential for developing a more thorough understanding of *E. coli*, its pathogenicity and the role played by associated plasmids. This study reports complete genomes for 15 well documented strains and a total of 34 plasmids associated with them, providing new data on the ComEC population carried by a healthy human adult. The diversity of ComEC is clearly illustrated by this dataset, with all phylogroups apart from C represented in Subject 24 over the almost two-year sampling period.

An important aspect of this study is that it provides information relating to two facets of successful gut colonisation, namely extended residency and dominance. Crucially, comparisons of the complete genome sequences generated here for isolates from their first and last appearances have unambiguously demonstrated that five ComEC strains from phylogroups A, B2 or D resided in Subject 24 for ≥20 days, with the longest residency ≥170 days. Strikingly, even the numbers and positions of insertion sequences in persistent strain chromosomes were conserved over the course of their residencies. These persistent strains were also dominant (defined as representing >80% of *E. coli* in a sample) in most samples (n=14/21) and two of them were never <95% of *E. coli* in the samples they appeared in (Figure 2). In contrast, only one of the ten strains that were isolated from just one sample (24.20-R5) was dominant (Figure 2). The availability of detailed gut residency data in conjunction with complete genome sequences will be critical in future comparative studies aiming to evaluate the differences between “successful” and “unsuccessful” ComEC. The complete genomes reported here add to those reported previously.^18^ Determination of sequences for the known persistent *E. coli* strains recovered in molecular studies^17,19,21,22^ would significantly increase this data set. In addition, genome sequences for strains that persisted in infants^60,61^ would be highly valuable in comparative studies.

The potential for horizontal gene transfer in the human gut has been appreciated for decades,^12–14^ but historic studies that examined gene transfer in the healthy human gut were limited in their ability to examine plasmids and distinguish bacterial hosts involved.^2^ Several studies presented evidence of transfer under selective pressure,^62–64^ or introduced donor and/or recipient strains,^13,14,65,66^ and in recent years examples of antibiotic resistance gene transfer in the guts of hospitalised patients have grown considerably.^67^ However, documented examples of transfer events occurring under natural conditions in the absence of direct selection for transconjugants remain scarce.^68^ The sampling protocol used here was designed to increase the chances of detecting natural transfer events, and yielded some examples. First, a Z-type antibiotic resistance plasmid transferred from one persistent strain to the next before undergoing a microevolution event in the recipient (Figure 3). Then, although the original donor was not detected, two strains acquired the same M-type arsenic resistance plasmid (Figure 4). These cases clearly represent plasmid transfer *in situ*.

This work highlights the value of an in-depth approach to the study of ComEC in healthy individuals. Identification and compilation of available genomes for ComEC strains with known residency characteristics from healthy humans living in the community will also advance the study of the relationship between ComEC and strains that have caused extraintestinal infections (ExPEC), expanding on the growing appreciation for the possible link between successful gut colonisation/persistence and infection proficiency.^22^

---

**Table S1:** Characteristics of Enterobacteriaceae isolated from Subject 24.

| **Sample** | **Study Day** | **Abundance Enterobacteriaceae (cfu/g)** | **Resistance phenotype** | **API20E profile** | **RAPD type** | **Phylo group** | **Strain** | **% *E. coli* in sample** |
|---|---|---|---|---|---|---|---|---|
| 1 | 1 | 2.2 x 10<sup>7</sup> | - | 5144572 | 1 | B2 | 24-S1 | 20.6 |
|  |  |  | - | 5144572 | 2 | B1 | 24.1-S2 | 16.8 |
|  |  |  | Ap CAZ CTX Tc | 5044552 | 3 | D | 24.1-R1 | 61.7 |
|  |  |  | ApCmSmSpSuTcTp | 5044542 | 4 | D | 24.1-R2 | 0.9<sup>1</sup> |
| 2 | 38 | 5.1 x 10<sup>7</sup> | - | 5144572 | 1 | B2 | 24-S1 | 68.4 |
|  |  |  | - | 5144572 | 5 | D | 24.2-S3 | 31.6 |
| 3 | 65 | 3.6 x 10<sup>7</sup> | - | 5044552 | 6 | B1 | 24.3-S4 | 78.0 |
|  |  |  | - | 5144572 | 1 | B2 | 24-S1 | 22.0 |
| 4 | 101 | 2.2 x 10<sup>7</sup> | - | 5144552 | 7 | B1 | 24.4-S5 | 77.6 |
|  |  |  | - | 5144572 | 1 | B2 | 24-S1 | 22.4 |
| 5 | 130 | 3.8 x 10<sup>7</sup> | - | 5144572 | 1 | B2 | 24-S1 | 100 |
| 6 | 171 | 2.9 x 10<sup>7</sup> | - | 5144572 | 1 | B2 | 24-S1 | 100 |
| 7 | 282 | 5.2 x 10<sup>7</sup> | - | 5144552 | 8 | B2 | 24-S6 | 100 |
| 8 | 318 | 1.4 x 10<sup>7</sup> | - | 5144552 | 8 | B2 | 24-S6 | 95.0 |
|  |  |  | - | 5144572 | 9 | B2 | 24.8-S7 | 5.0 |
| 9 | 346 | 1.1 x 10<sup>7</sup> | - | 5144552 | 8 | B2 | 24-S6 | 100 |
| 10 | 374 | 1.9 x 10<sup>7</sup> | - | 5144552 | 8 | B2 | 24-S6 | 100 |
| 11 | 402 | 2.4 x 10<sup>7</sup> | - | 5144552 | 8 | B2 | 24-S6 | 100 |
| 12 | 437 | 1.2 x 10<sup>7</sup> | ApSmSuTp | 1044452 | 10 | A | 24-R3 | 100 |
| 13 | 469 | 2.5 x 10<sup>7</sup> | ApSmSuTp | 1044452 | 10 | A | 24-R3 | 100 |
| 14 | 493 | 7.3 x 10<sup>6</sup> | ApSmSuTp | 1044452 | 10 | A | 24-R3 | 100 |
| 15 | 520 | 2.9 x 10<sup>7</sup> | ApSmSuTp | 1044452 | 10 | A | 24-R3 | 100 |
| 16 | 548 | 1.3 x 10<sup>7</sup> | - | 5044572 | 11 | A | 24-S8 | 55.0 |
|  |  |  | ApSmSuTp | 5044572 | 11 | A | 24-S8R | 45.0 |
| 17 | 588 | 5.7 x 10<sup>7</sup> | - | 5044572 | 11 | A | 24-S8 | 80.0 |
|  |  |  | - | 5144572 | 12 | A | 24.17-S9 | 20.0 |
| 18 | 609 | 2.8 x 10<sup>7</sup> | - | 5044572 | 11 | A | 24-S8 | 100 |
| 19 | 647 | 5.3 x 10<sup>7</sup> | - | 5144562 | 13 | A | 24.19-S10 | 55.0 |
|  |  |  | - | 5044572 | 11 | A | 24-S8M | 45.0 |
| 20 | 680 | 1.2 x 10<sup>7</sup> | Tc | 5044512 | 14 | A | 24.20-R5 | 99.0 |
|  |  |  | - | 5144572 | 15 | D | 24-S11 | 1.0 |
| 21 | 700 | 5.8 x 10<sup>7</sup> | - | 5144572 | 15 | D | 24-S11M | 100 |
|  |  |  | - | 3204772 | 16 | - | 24.21-Cfr | -<sup>1</sup> |
|  |  |  | - | 5215773 | 17 | - | 24.21-Kva | -<sup>2</sup> |

<sup>1</sup> Estimated by calculating the abundance of colonies unique to this phenotype that grew on antibiotic-containing agar plates relative to the total *E. coli* abundance calculated from non-selective MacConkey agar plates.
<sup>2</sup> 27/100 presumptive *E. coli* colonies picked from Sample 21 grew on Simmons Citrate Agar and were determined to be the same strain using RAPD PCR. Identified as *Citrobacter freundii* using API20E, confirmed with its complete genome sequence.
<sup>2</sup> Detected as large mucoid colonies that were clearly distinct from presumptive *E. coli*. Estimated to account for approximately 5% of Enterobacteriaceae in Sample 21. Originally identified as *Klebsiella pneumoniae* using API20E, complete genome sequence revealed species identity *Klebsiella variicola*.

**Table S2:** Changes in the chromosomes of Subject 24 persistent E. coli strains.

| Strain | Change<sup>1</sup> | Position | ORF | AA change | Notes |
|---|---|---|---|---|---|
| **24-S1** | C > T | 75,120 | *spoT* | A397T | mediator of stringent response; degrades ppGpp > GDP |
|  | C > T | 810,290 | - | - | 372 bp upstream of *kpsMII* |
|  | C > T | 1,088,367 | *rpoS* | Q71* | alternative sigma factor (stress response) |
|  | CTGG > ---- | 1,781,374 | *wcaA* | P100V; G101* | glycosyltransferase for colanic acid synthesis |
|  | T > - | 4,613,509 | - | - | 23S ribosomal rRNA |
|  | G > A | 4,732,631 | - | - | between 16S and 23S rRNA genes |
| **24-S6** | T > A | 103,950 | *waaL* | L22* | ligase |
|  | 12 bp Δ | 3,796,694 | *rpnA* | 4 AA shorter | putative transposase |
| **24-R3** | C > T | 1,166,859 | - | - | 11 bp upstream of *emrR* (negative regulator of *emrAB* efflux genes) |
|  | C > A | 1,345,631 | *purM* | G114C | purine biosynthesis pathway |
|  | G > A | 1,633,835 | - | - | downstream of two oppositely oriented ORFs |
| **24-S8** | C > T | 813,002 | *kpsS* | R204H | capsular polysaccharide modification protein |
|  | A > T | 2,526,023 | orf | Y2* | superinfection exclusion B family protein |
|  | G > A | 2,686,942 | *misL?* | A632T | outer membrane autotransporter barrel |
| **24-S11** | A > - | 1,880,326 | *wbbD* | frameshift | glycosyltransferase |
|  | ---- > TATA | 1,880,906 | *wbbD* | frameshift |  |

<sup>1</sup> The sequence in the first appearance isolate’s chromosome is given first, or only the change seen in the final appearance is listed.

**Figure S1:**
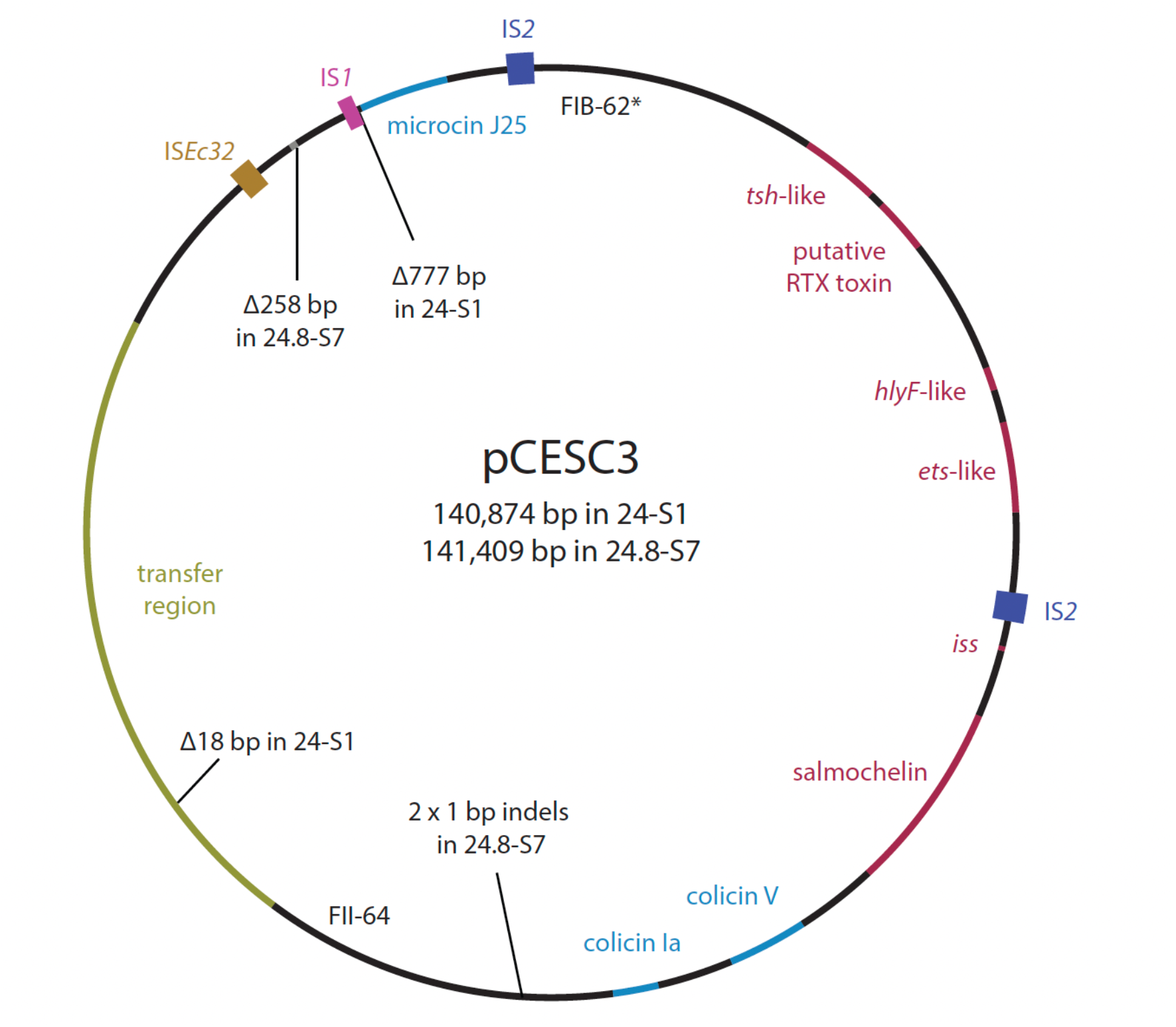
Basic circular map of pCESC3. Regions of the plasmid that encode bacteriocin (blue), conjugative transfer (yellow), and ‘virulence’ (red) determinants are highlighted and labelled. The *tsh-, hlyF-,* and ete-like genes are related, but not identical to, their equivalents in classical ColV plasmids. Insefrtion sequences are shown as labelled boxes. Differences that account for the size discrepancy between pCESC3 in 24-Sl and pCESC2-v2 in 24.8-S7 are labelled. The FIB-62* replicon differs from the FIB-62 reference sequence in PubMLST by a single base change.

## Notes

### Competing Interest Statement

The authors have declared no competing interest.

### Funding

- Australian National Health and Medical Research Council: GNT1194978
- Department of Education: Australian Postgraduate Award (2013 - 2016)

